# Golgi structural defect impairs glycosaminoglycan synthesis, sulfation, and secretion

**DOI:** 10.1101/2021.10.14.464461

**Authors:** Erpan Ahat, Yuefan Song, Ke Xia, Whitney Reid, Jie Li, Sarah Bui, Fuming Zhang, Robert J Linhardt, Yanzhuang Wang

## Abstract

Synthesis of glycosaminoglycans such as heparan sulfate (HS) and chondroitin sulfate (CS) occurs in the lumen of the Golgi but the relationship between Golgi structural integrity and glycosaminoglycan synthesis is not clear. In this study, we disrupted the Golgi structure by knocking out GRASP55 and GRASP65 and determined its effect on the synthesis, sulfation, and secretion of HS and CS. We found that GRASP depletion increased HS synthesis while decreasing CS synthesis in cells, altered HS and CS sulfation, and reduced both HS and CS secretion. Using proteomics, RNA-seq and biochemical approaches, we identified EXTL3, a key enzyme in the HS synthesis pathway, whose level is upregulated in GRASP knockout cells; while GalNacT1, an essential CS synthesis enzyme, is robustly reduced. In addition, we found that GRASP depletion decreased HS sulfation via the reduction of PAPSS2, a bifunctional enzyme in HS sulfation. Our study provides the first evidence that Golgi structural defect may significantly alter the synthesis and secretion of glycosaminoglycans.

## Introduction

The Golgi apparatus is a central station in the intracellular trafficking pathway and serves as the principal hub for sorting and post-translational modifications of proteins and lipids (Li et al., 2019). The basic structure of the Golgi is a stack of flattened cisternae. In mammalian cells, multiple Golgi stacks are latterly linked into a ribbon-like structure located in the perinuclear region of the cell. It has been previously demonstrated that two Golgi peripheral membrane proteins, GRASP55 and GRASP65, function as the “glue” that links Golgi membranes together and facilitates Golgi stacking and ribbon formation. Knocking down or knocking out either one of these two GRASP proteins decreases the number of cisternae per stack, whereas depleting both GRASPs disrupts the entire Golgi structure (Xiang and Wang, 2010, Bekier et al., 2017, Ahat et al., 2019a).

Functional studies revealed that destruction of the Golgi structure by GRASP-depletion accelerates protein trafficking in the Golgi, but impairs accurate N-glycosylation and protein sorting (Xiang et al., 2013). At the cellular level, GRASP depletion reduces cancer cell attachment and invasion mainly through the reduction of α5β1 integrin synthesis (Ahat et al., 2019b), indicating a role of GRASPs and/or the Golgi structure in transcription regulation. In addition to Golgi structure formation, GRASPs are also involved in autophagy and unconventional secretion. Under starvation or stress conditions, GRASP55, but not GRASP65, translocates from the Golgi to other membrane structures such as autophagosomes and endoplasmic reticulum (ER) to regulate autophagy and unconventional secretion of certain cytosolic or transmembrane proteins (Zhang et al., 2018, Zhang and Wang, 2018, Gee et al., 2011, Cruz-Garcia et al., 2017, Nüchel et al., 2021, Ahat et al., 2021).

Glycosaminoglycans (GAGs) are main components of the cell surface glycome and extracellular matrix (Annaval et al., 2020). GAGs are long linear polysaccharides consisting of repeating disaccharide units. Based on the core disaccharide structures, GAGs are classified into three major forms, heparan sulfate (HS), chondroitin sulfate (CS), and hyaluronan (HA). While HA is synthesized at the plasma membrane and released, HS and CS are synthesized and attached to serine residues of cargo proteins in the Golgi, from where they are transported to the cell surface and secreted to the extracellular space. The biosynthesis of both HS and CS begins with the formation of a tetrasaccharide linker on a serine residue in a protein core, which is subsequently diversified to HS or CS depending on the subsequent enzymatic reactions (Figure 1A) (Annaval et al., 2020).

**Figure 1.**
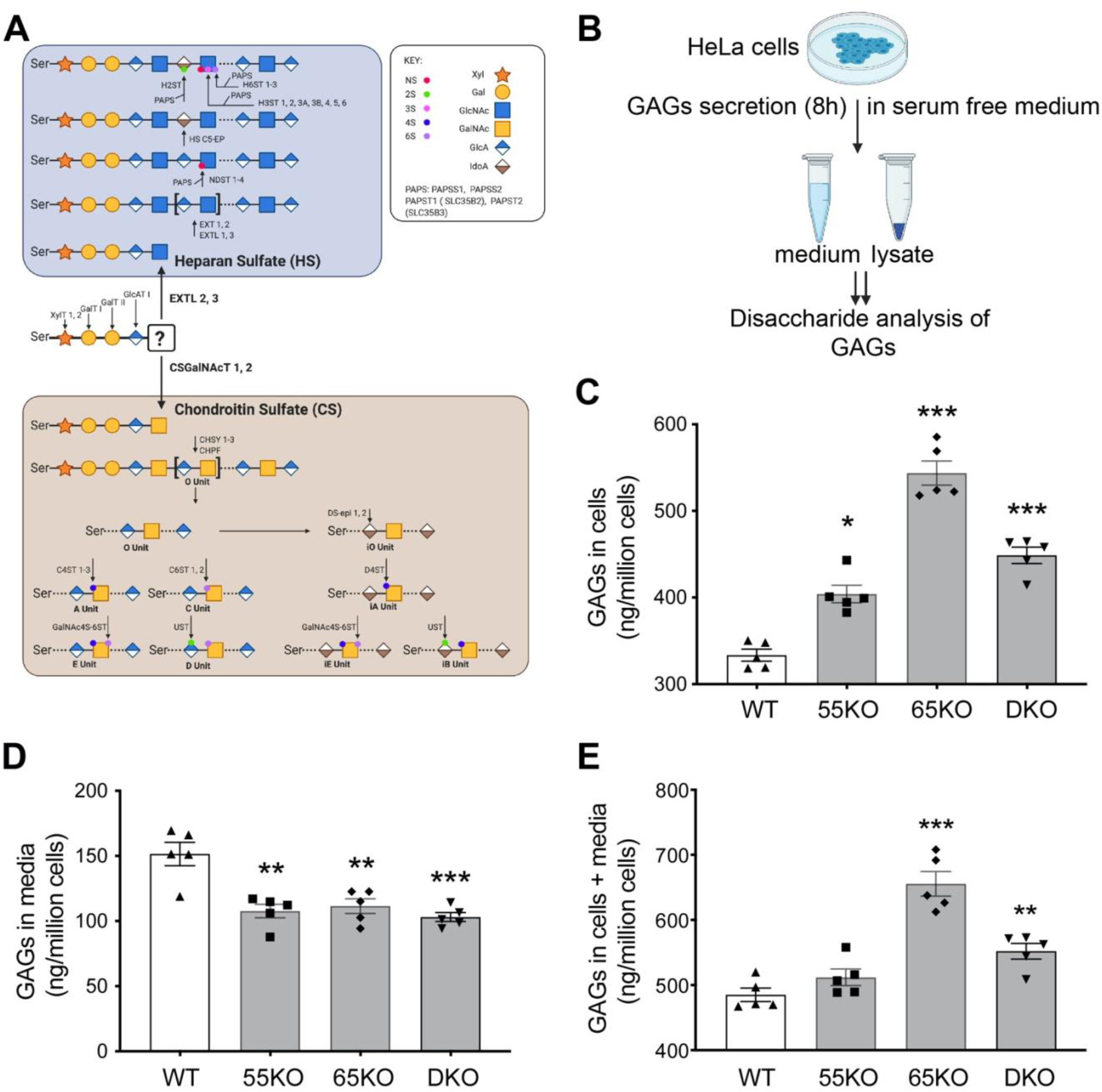
Golgi structure disruption by GRASP KO increases GAG synthesis but reduces its secretion. **(A)** Schematic diagram illustrating the HS and CS synthesis pathways; major enzymes in each pathway are indicated at their designated steps of reactions. **(B)** Schematic workflow for cell lysate and medium sample preparation to analyze GAGs by LC-MS. **(C)** GRASP KO increases the amount of GAGs in cells. Shown are the amounts of GAGs in the lysates of wildtype (WT), GRASP55 knockout (55KO), GRASP65 knockout (65KO), and GRASP55 and GRASP65 double knockout (DKO) HeLa cells. **(D)** GRASP KO decreases GAG secretion. WT and indicated GRASP KO cells were incubated in serum-free medium for 8 h and GAGs in the conditioned media were analyzed by LC-MS. **(E)** GRASP KO increases the total amount of GAGs in cells and media. Shown are the total amount of GAGs per million cells in both cell lysates and conditioned medium in each cell line. Results are presented as mean ± SEM, statistical analysis was assessed by comparing KO cells to WT cells by student’s t-test. *, p<0.05; **, p<0.01; ***, p<0.001.

For HS, the initiation of HS biosynthesis occurs as the transferases Exostosin-like 2 (EXTL2) and 3 (EXTL3) transfer an *N*-Acetylglucosamine (GlcNAc) to the initial linker chain (Figure 1A) (Kim et al., 2001, Kitagawa et al., 1999). Next, Exostosin-1 (EXT1) and −2 (EXT2), and Exostosin-like 1 (EXTL1) and EXTL3, extend the chain by alternatively transferring GlcNAc and D-Glucuronic acid (GlcA) residues to the sugar chain (Kreuger and Kjellen, 2012). This HS chain is then modified by the enzymes bifunctional heparan sulfate *N*-deacetylase/*N*-sulfotransferase 1-4 (NDST1-4) that have both N-deacetylase and N-sulfotransferase activities (Carlsson et al., 2008). Other enzymes involved in HS sulfation include heparan sulfate 2-*O*-sulfotransferase (H2ST), 6-*O*-sulfotransferases (H6ST1-3), and 3-*O*-sulfotransferases (H3ST) 1, 2, 3A, 3B, 4, 5, and 6 (Maeda, 2015). These NDST enzymes and sulfotransferases rely on sulfur donor bifunctional 3’-phosphoadenosine 5’-phosphosulfate synthase (PAPS) which are synthesized by 3’-phosphoadenosine 5’-phosphosulfate synthase-1 (PAPSS1) and −2 (PAPSS2) in the cytosol and transported by adenosine 3’-phospho 5’-phosphosulfate transporter 1 (PAPST1) and PAPST2 into the Golgi lumen (Fuda et al., 2002, Stelzer et al., 2007, Kamiyama et al., 2006, Dick et al., 2015). Most HS chains are sulfated, which significantly affects their activity and function. For example, it has been shown that 3-*O*-sulfation of HS increases its binding with Tau at the cell surface, which facilitates Tau internalization (Zhao et al., 2020).

For CS, chain formation initiates with *N*-acetylgalactosaminyltransferases, GalNAcT1-2, and is then elongated by chondroitin sulfate synthase 1-3 (CHSY1-3) and chondroitin sulfate glucuronyltransferase (CHPF) that alternatively transfer GalNAc and GlcA residues to the sugar chain (Figure 1A) (Maeda, 2015). CS is sulfated by sulfotransferases chondroitin 4-sulfotransferase 1-3 (C4ST1-3), chondroitin 6-sulfotransferase 1-2 (C6ST1-2) and *N*-acetylgalactosamine 4-sulfate 6-*O*-sulfotransferase (GalNAc4S-6ST), or by dermatan sulfate epimerases 1-2 (DS-epi 1-2) if the 2-*O*-sulfated D-glucuronic acid (GlcA) residues are C5-epimerized to L-iduronic acid (IdoA). Subsequently, when the chain is further sulfated by dermatan sulfotransferase D4ST, uronyl 2-*O*-sulfotransferase (UST), or GalNAc4S-6ST, it is referred to as dermatan sulfate (DS) (Maeda, 2015).

While the synthesis of the tetrasaccharide linker of HS and CS is initiated in the ER or ERGIC, the bulk parts of HS and CS are synthesized in the Golgi lumen (Prydz, 2015), similar to that of *N-*linked glycans. Consistently, most enzymes involved in HS and CS synthesis, such as EXT1 and EXT2 in HS synthesis and GalNAcT-1 in CS synthesis, reside in the Golgi (Kobayashi et al., 2000, Uliana et al., 2006). Therefore, it is reasonable to speculate that Golgi structural defect may significantly impact the synthesis of HS and CS as *N*-glycans as we previously showed (Xiang et al., 2013). Consistently, depletion of certain subunits of the Conserved Oligomeric Golgi (COG) complex, which transports Golgi enzymes to their proper locations within the Golgi, reduces GAG modification (Adusumalli et al., 2021); while depletion of giantin, another membrane tether in the Golgi, reduces the mRNA level of polypeptide *N*-acetylgalactosaminyltransferase 3 (GALNT3) (Stevenson et al., 2017). However, the relationship between Golgi structural integrity and GAG synthesis has not been systematically explored.

In this study, we disrupted the Golgi structure by knocking out GRASP55 and GRASP65 and determined the effect on GAG synthesis, sulfation, and secretion. We also performed proteomic and RNA-seq analysis to identify the enzymes whose alternation is responsible for the defects in HS and CS synthesis in GRASP knockout (KO) cells.

## Results

### GRASP KO increases GAG synthesis but decreases their secretion

Given that GRASP55 and GRASP65 are major regulators of Golgi stack formation, we knocked them out, single or in combination, in HeLa cells to disrupt the Golgi structure (Bekier et al., 2017), and thereby determined the effect on GAG synthesis. As shown in Figure 1B, we cultured wildtype (WT), GRASP55 knockout (55KO), GRASP65 knockout (65KO), and GRASP55 and GRASP65 double knockout (DKO) cells in serum-free medium for 8 h, collected cell lysate and conditioned media, and performed GAG analysis by liquid chromatography-mass spectrometry (LC-MS) (Supplemental Tables 1 and 2). The amount of GAGs, including (HS, CS and HA) in the cell lysate was increased in GRASP KO cells, with its level in 65KO the highest (Figure 1C). In contrast to the cell lysate, the amount of GAGs in the conditioned media was reduced (Figure 1D). The total amount of GAGs (cells + media) was higher in GRASP KO cells compared to WT, again with 65KO to be the highest (Figure 1E). In summary, disruption of the Golgi structure by GRASP depletion increases GAG synthesis while reducing its secretion.

### GRASP KO increases HS synthesis but decreases its sulfation and secretion

Given that HS and CS but not HA are synthesized in the Golgi, we further characterized HS and CS synthesis and sulfation in GRASP KO cells. When the Golgi structure was disrupted by GRASP KO, HS synthesis was significantly increased compared to WT cells as analyzed by LC-MS (Figure 2A; Supplemental Table 1–2). The increase of HS in GRASP KO cells or at the cell surface was confirmed by immunostaining of HS with an anti-HS antibody 10E4 followed by immunofluorescence microscopy with or without permeabilization (Figure 2B; Figure S1A). The HS signal by this antibody was specific as it was largely quenched by preincubation of the antibody with heparin (Figure S1B). The increased level of HS in GRASP KO cells was further validated by flow cytometry (Figure 2C).

**Figure 2.**
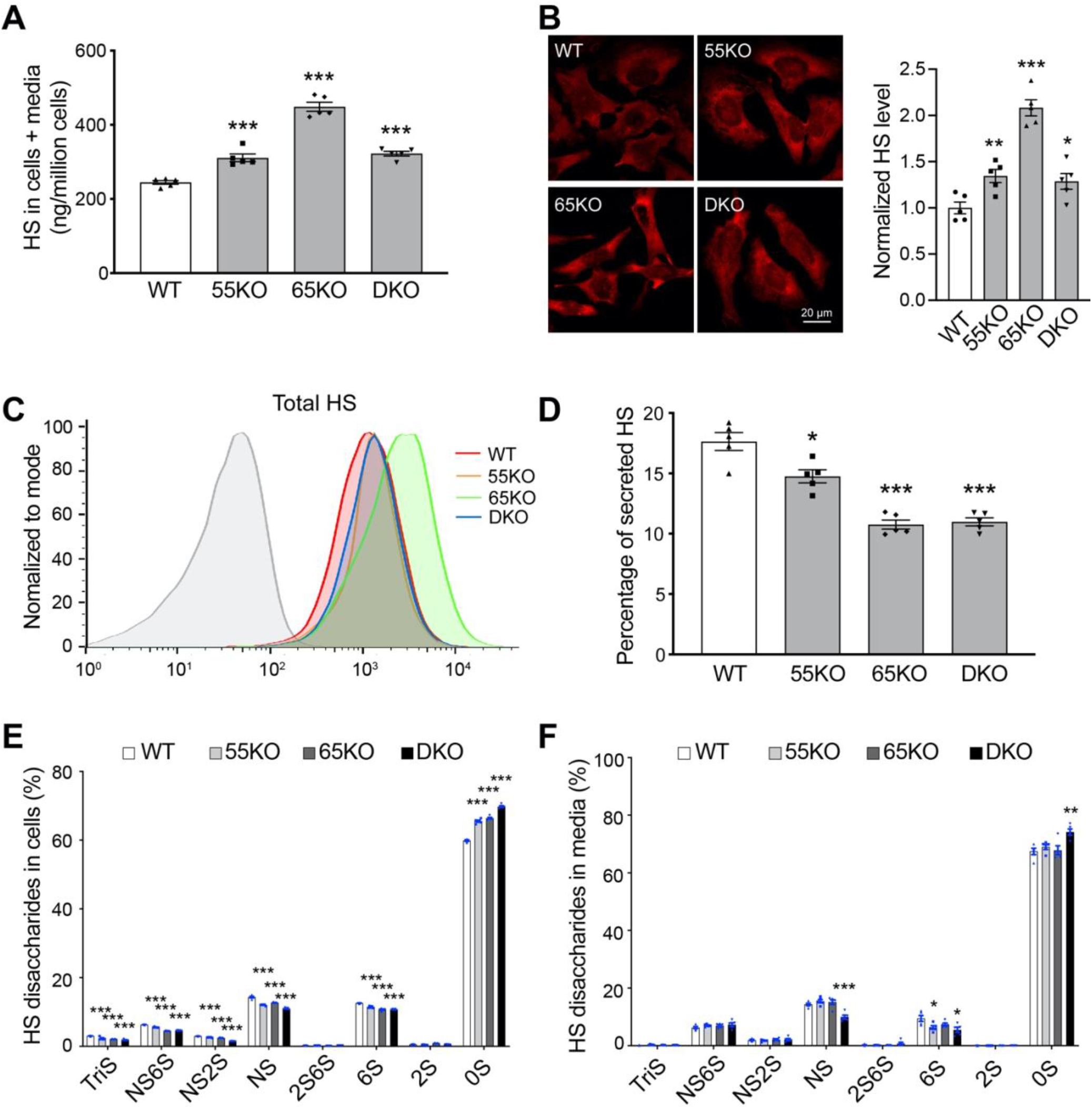
GRASP KO increases HS synthesis but reduces its sulfation and secretion. **(A)** GRASP KO increases HS synthesis analyzed by LC-MS. Shown are the total amount of HS in both cells and medium of indicated cell lines. **(B)** GRASP KO increases HS synthesis analyzed by immunofluorescence microscopy. Indicated cells were permeabilized and stained for HS with an HS antibody 10E4. Shown are microscopic images (left) and quantitation (right). **(C)** GRASP KO increases HS synthesis analyzed by flow cytometry. Indicated cells were permeabilized, stained for HS with an HS antibody and analyzed by flow cytometry. **(D)** GRASP KO decreases HS secretion. The percentage of secreted HS (HS in media/HS in cell lysate and conditioned media) was analyzed by LC-MS. **(E)** GRASP KO reduces HS sulfation in the cell lysate. Shown are the percentage of each sulfated form of HS in cells. **(F)** GRASP KO reduces HS sulfation in the conditioned media. Shown are the percentage of each sulfated form of HS in the conditioned media.

HS is covalently linked to core proteins that are secreted by cells, so we analyzed the level of HS in the conditioned media. GRASP KO largely reduced HS secretion compared to that of WT cells (Figure 2D). Although the absolute amount of HS in the media was not reduced by GRASP KO, the percentage of HS in the media was significantly lower in the KOs than WT due to the increased HS synthesis (Supplemental Tables 1–2). Given the importance of HS sulfation, we also quantified the different sulfated forms of HS in cells and conditioned media by 2-aminoacridone (AMAC) labeling and LC-MS (Supplemental Tables 3 and 4). In contrast to the increased level of HS synthesis, the overall sulfation of HS was significantly reduced by GRASP KO in both the cells and conditioned media (Figure 2E-F). This indicates that Golgi destruction via GRASP depletion negatively regulates the sulfation pathway of HS. Taken together, disruption of the Golgi structure by GRASP KO increases HS synthesis but decreases its sulfation and secretion.

### GRASP KO decreases CS synthesis and secretion

Like HS, CS is synthesized in the Golgi and sulfated. HS and CS share the same tetrasaccharide precursor, which branches into either the HS pathway via the action of the EXTL enzymes, or the CS pathway via the reaction of the GalNAcT enzymes (Figure 1A). Therefore, it is reasonable to speculate that increased branching into the HS pathway may lead to reduced branching into the CS pathway. Indeed, GRASP KO reduced CS synthesis as analyzed by LC-MS, the opposite to HS (Figure 3A; Supplemental Tables 1 and 2). To confirm this result by an alternative approach, we stained WT and GRASP KO cells with an CS antibody (CS-56) and analyzed the levels of CS by fluorescence microscopy and flow cytometry. Consistent with the LC-MS results, GRASP KO reduced the level of CS in cells compared to WT (Figure 3B-C). Here, the degree of CS reduction examined by microscopy and flow cytometry was more dramatic than by LC-MS. The cause of this difference could be that LC-MS includes both CS and DS in the results, while the CS antibody only recognizes CS but not DS (Avnur and Geiger, 1984), which more accurately reflects the CS level in cells.

**Figure 3.**
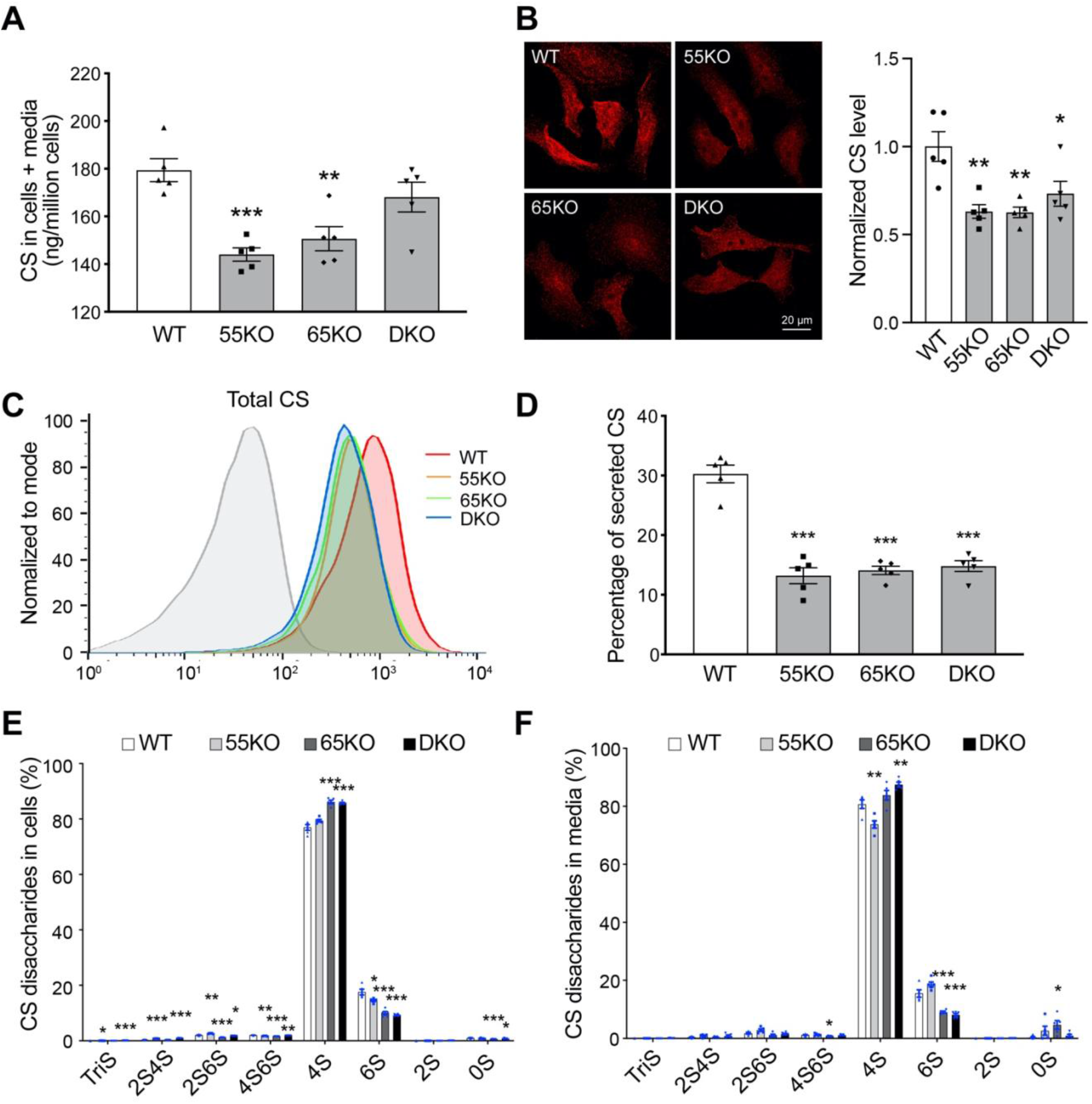
GRASP KO reduces CS synthesis and secretion. **(A)** GRASP KO reduces CS synthesis analyzed by LC-MS. Shown are the total amount of CS in both cells and medium of indicated cell lines. **(B)** GRASP KO decreases CS synthesis analyzed by immunofluorescence microscopy. Indicated cells were permeabilized and stained for CS with a CS antibody (CS-56). Shown are microscopic images (left) and quantitation (right). **(C)** GRASP KO decreases CS synthesis shown by flow cytometry. Indicated cells were permeabilized, stained for CS with a CS antibody and analyzed by flow cytometry. **(D)** GRASP KO decreases CS secretion. The percentage of secreted CS (CS in media/HS in cell lysate and conditioned media) was analyzed by LC-MS. **(E)** GRASP KO alters CS sulfation in the cell lysate. Shown are the percentage of each sulfated form of CS in cells. **(F)** GRASP KO alters CS sulfation in the conditioned media. Shown are the percentage of each sulfated form of CS in the conditioned media. Note that GRASP depletion increased 4-sulfation while decreases 6-sulfation in both the cell lysate (E) and conditioned media (F).

Next, we analyzed the secretion of CS in WT and GRASP KO cells. Both the amount and percentage of CS in the conditioned media were largely reduced by GRASP KO (Figure 3D; Supplemental Tables 1–2). Lastly, we analyzed the different subtypes of CS sulfation and found that GRASP KO increased 4-sulfation but decreased 6-sulfation in both the cell lysate and conditioned media (Figure 3E-F; Supplemental Tables 5 and 6). Taken together, disruption of the Golgi structure by GRASP KO decreases CS synthesis, alters its sulfation, and decreases its secretion.

### GRASP KO regulates key enzymes in HS and CS synthesis and sulfation

The regulation of HS and CS synthesis and sulfation is complex and involves numerous enzymes (Figure 1A). A majority of these enzymes are localized in the Golgi and thus their level and localization could be regulated by Golgi structural changes. Therefore, we performed systematic RNA-seq and proteomic analysis of WT and GRASP KO cells to identify genes related to the observed alterations in HS and CS synthesis and sulfation. As expected, many enzymes involved in HS and CS synthesis were affected by GRASP KO (Table 1). Consistent with the increased HS level in cells, the mRNA levels of several HS synthesis enzymes, such as EXTL2, EXTL3, EXT1, NDST1 and Adenosine 3’-phospho 5’-phosphosulfate transporter 3 (SLC35B3), were increased in GRASP KO compared to WT cells. Similar to the trend of CS reduction in GRASP KO cells, the mRNA levels of many CS synthesis enzymes, including GalNAcT1, CHSY1, C4ST2, GalNAc4S-6ST, C6ST1 and DS-epi1, were all decreased upon GRASP KO (Figure 4A). The altered expression of HS and CS synthesis enzymes in GRASP KO cells was further confirmed at the protein level by proteomic analysis (Table 1) and Western blots (Figure 4B). GRASP KO increased the protein level of EXTL3, a key enzyme in HS synthesis, while decreased the level of GalNAcT1 in the CS synthesis pathway (Figure 4B-C). In addition to EXTL3, the protein levels of EXT1 and EXT2 were slightly higher in 65KO cells, which might explain why 65KO cells have the highest HS level in all cell lines.

**Table 1.** GRASP KO alters HS and CS synthesis enzymes at both the mRNA and protein levels. WT and GRASP KO cells were analyzed by proteomic and RNA-seq analyses. Shown are the fold changes of the indicated GAG synthesis enzymes in GRASP KO cells compared to that in WT cells. Enzymes in bold were further tested by Western blot. NA, data not available; ND, not detected.

| **Enzymes** | **Proteomics** |  |  |  | **RNA-seq** |  |  |  |  |  |
| --- | --- | --- | --- | --- | --- | --- | --- | --- | --- | --- |
|  | **55KO/WT** | **p-value** | **65KO/WT** | **p-value** | **log2 (55KO/WT)** | **p-value** | **log2 (65KO/WT)** | **p-value** | **log2 (DKO/WT)** | **p-value** |
| **<u>Common Enzymes</u>** |  |  |  |  |  |  |  |  |  |  |
| XylT1 | ND |  | ND |  | -0.009 | NA | 6.511E-04 | NA | -0.007 | NA |
| XylT2 | ND |  | ND |  | 0.100 | 0.770 | -0.250 | 0.290 | -0.100 | 0.740 |
| GalT1 (B4GALT7) | ND |  | ND |  | 0.210 | 0.440 | -0.170 | 0.500 | 0.140 | 0.590 |
| GalT2 | ND |  | ND |  | -0.850 | 0.020 | -0.410 | 0.340 | -0.110 | 0.840 |
| GlcAT1 (B3GAT3) | ND |  | ND |  | -0.370 | 0.210 | -0.110 | 0.750 | 0.020 | 0.960 |
| **<u>Enzymes in HS synthesis</u>** |  |  |  |  |  |  |  |  |  |  |
| EXTL2 | ND |  | ND |  | -0.170 | 0.420 | 0.320 | 0.030 | 0.090 | 0.670 |
| **EXTL3** | ND |  | ND |  | 0.330 | 0.120 | 0.130 | 0.620 | 0.290 | 0.160 |
| EXTL1 | ND |  | ND |  | -0.010 | NA | -0.010 | NA | -0.010 | NA |
| **EXT1** | ND |  | ND |  | 0.200 | 0.420 | 0.660 | 3.090E-05 | 0.180 | 0.430 |
| **EXT2** | ND |  | ND |  | -0.080 | 0.790 | -0.100 | 0.690 | -0.230 | 0.240 |
| NDST 1 | ND |  | ND |  | 0.230 | 0.260 | 0.120 | 0.600 | -0.170 | 0.400 |
| NDST 2 | ND |  | ND |  | 0.280 | 0.620 | -0.030 | 0.960 | -0.050 | 0.940 |
| NDST 3 | ND |  | ND |  | -0.210 | 0.700 | -0.460 | 0.200 | 0.100 | 0.850 |
| NDST 4 | ND |  | ND |  | ND |  | ND |  | ND |  |
| PAPSS 1 | 0.047 | 0.411 | -0.093 | 0.082 | 3.733E-03 | 0.990 | -0.060 | 0.790 | -0.120 | 0.460 |
| **PAPSS 2** | -0.825 | 1.854E-05 | -0.873 | 2.289E-06 | -0.910 | 7.920E-10 | -0.690 | 2.269E-06 | -0.690 | 3.599E-06 |
| PAPST 1 (SLC35B2) | -0.141 | 0.034 | -0.069 | 0.256 | -0.120 | 0.750 | 0.380 | 0.080 | 0.400 | 0.060 |
| PAPST 2 (SLC35B3) | ND |  | ND |  | 0.290 | 0.190 | 0.070 | 0.800 | 0.370 | 0.050 |
| HS C5-EP (HSGLCE) | ND |  | ND |  | ND |  | ND |  | ND |  |
| H2ST | ND |  | ND |  | ND |  | ND |  | ND |  |
| H3ST 1 | ND |  | ND |  | ND |  | ND |  | ND |  |
| H3ST 2 | ND |  | ND |  | ND |  | ND |  | ND |  |
| H3ST 3A | ND |  | ND |  | ND |  | ND |  | ND |  |
| H3ST 3B | ND |  | ND |  | ND |  | ND |  | ND |  |
| H3ST 4 | ND |  | ND |  | ND |  | ND |  | ND |  |
| H3ST 5 | ND |  | ND |  | ND |  | ND |  | ND |  |
| H3ST 6 | ND |  | ND |  | ND |  | ND |  | ND |  |
| H6ST 1 | ND |  | ND |  | ND |  | ND |  | ND |  |
| H6ST 2 | ND |  | ND |  | ND |  | ND |  | ND |  |
| H6ST 3 | ND |  | ND |  | ND |  | ND |  | ND |  |
| **<u>Enzymes in CS synthesis</u>** |  |  |  |  |  |  |  |  |  |  |
| **GalNAcT1 (GALNT1)** | -0.404 | 8.206E-05 | -0.521 | 1.34308E-05 | -0.521 | 2.856E-09 | -0.884 | 3.668E-26 | -1.135 | 7.254E-42 |
| GalNAcT2 (GALNT2) | -0.095 | 0.094 | 0.096 | 0.111 | -0.140 | 0.480 | 0.290 | 0.040 | -0.290 | 0.040 |
| CHPF2 | 0.159 | 0.368 | 0.197 | 0.227 | 0.040 | 0.950 | -0.050 | 0.910 | 0.200 | 0.540 |
| CHSY1 | ND |  | ND |  | -0.880 | 8.023E-07 | -0.693 | 9.417E-05 | -0.579 | 1.442E-03 |
| CHSY2 | ND |  | ND |  | NA |  | NA |  | NA |  |
| CHSY3 | ND |  | ND |  | -0.110 | 0.790 | 0.050 | 0.900 | 0.110 | 0.740 |
| C4ST 1 (CHST11) | ND |  | ND |  | -0.184 | 0.472 | 0.163 | 0.461 | -0.177 | 0.424 |
| C4ST 2 (CHST12) | ND |  | ND |  | -0.092 | 0.732 | -0.022 | 0.935 | -0.026 | 0.925 |
| C4ST 3 (CHST13) | ND |  | ND |  | -0.270 | 0.635 | 0.480 | 0.240 | 0.903 | 0.008 |
| GalNAc4S-6ST (CHST15) | ND |  | ND |  | -0.083 | 0.866 | -0.734 | 0.003 | -0.678 | 0.007 |
| C6ST 1 (CHST3) | ND |  | ND |  | -0.242 | 0.471 | -0.641 | 0.005 | -0.839 | 1.600E-04 |
| C6ST 2 (CHST4) | ND |  | ND |  | 0 | NA | 0 | NA | 0.013 | NA |
| UST | ND |  | ND |  | ND |  | ND |  | ND |  |
| DS-epi 1 (DSE) | ND |  | ND |  | -0.715 | 6.146E-10 | -1.274 | 3.265E-28 | -0.788 | 7.330E-12 |
| DS-epi 2 (DSEL) | ND |  | ND |  | 0.721 | 8.811E-05 | 0.505 | 8.939E-03 | -0.001 | 9.967E-01 |
| D4ST | ND |  | ND |  | ND |  | ND |  | ND |  |
| **<u>Enzymes in HA synthesis</u>** |  |  |  |  |  |  |  |  |  |  |
| HAS1 | ND |  | ND |  | -0.009 | NA | -0.009 | NA | -0.007 | NA |
| HAS2 | ND |  | ND |  | 0.131 | 0.850 | 0.088 | 0.888 | 0.390 | 0.356 |
| HAS3 | ND |  | ND |  | 0.009 | 0.992 | -0.132 | 0.819 | 0.066 | 0.915 |

**Figure 4.**
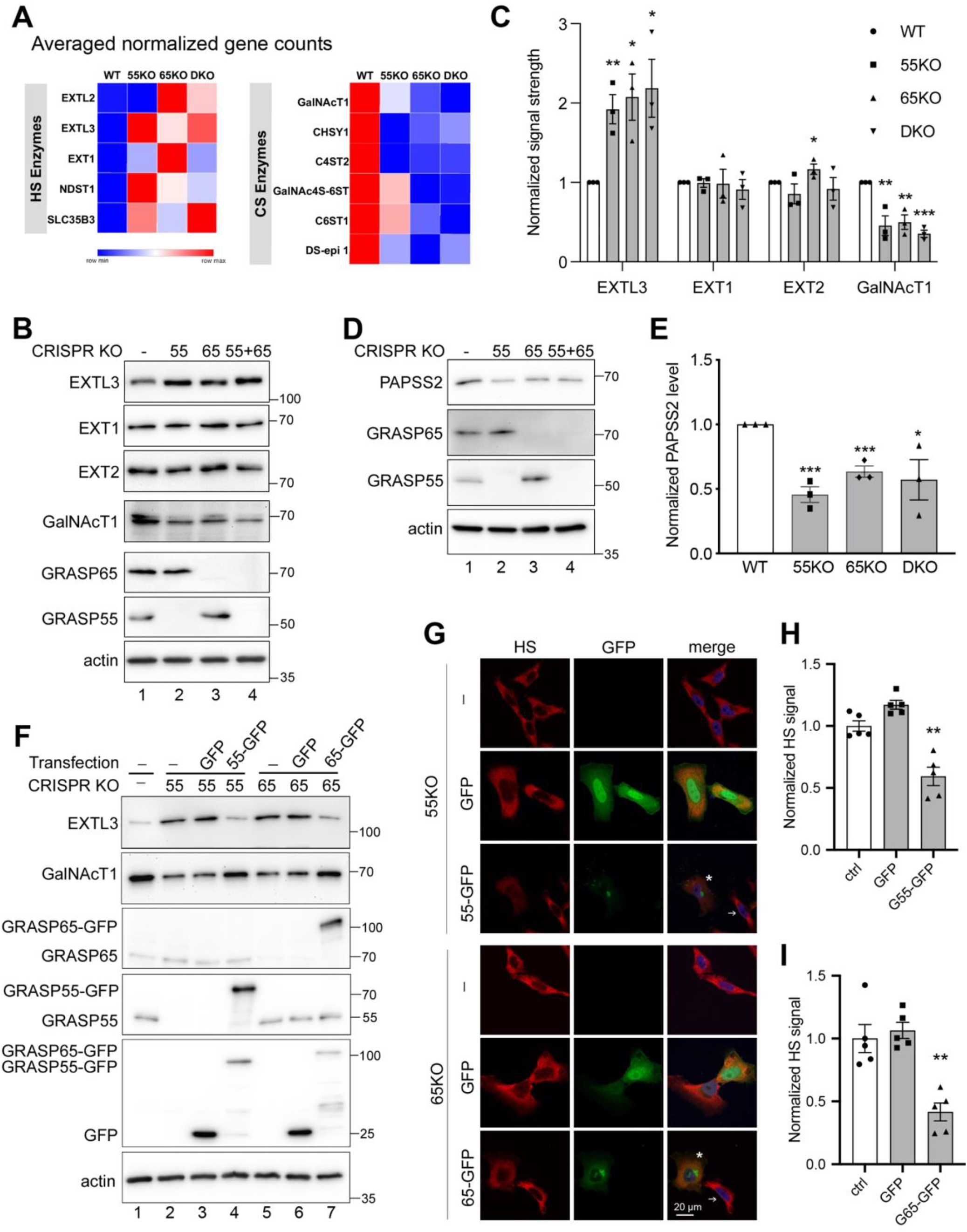
GRASP KO alters the expression level of GAG synthesis and sulfation enzymes. **(A)** GRASP KO increases the expression of HS synthesis enzymes while decreases CS synthesis enzymes. Results are based on RNA-Seq analysis of each cell line for indicated genes. **(B)** Quantification of C. **(C)** GRASP KO increases the protein level of HS synthesis enzymes while decreases that of CS synthesis enzymes. Cell lysate of indicated cells were analyzed for three HS synthesis enzymes EXTL3, EXT1 and EXT2, and a key CS synthesis enzyme GalNAcT1. Note the increased level of EXTL3 and decreased GalNAcT1 level in GRASP KO cells. Results are representative of three independent experiments. **(D)** GRASP KO decreases the protein level of the HS sulfation enzyme PAPSS2. Shown are representative Western blots of indicated proteins in the four cell lines from three independent experiments. **(E)** Quantitation of D. **(F)** Re-expression of GRASP proteins in GRASP KO cells corrects the expression level of HS and CS synthesis enzymes. Indicated cell lines were transfected with GRASP constructs and probed for EXTL3, GalNAcT1, GRASP65, GRASP55, GFP, and actin. The major enzymes EXTL3 and GalNAcT1 in 55KO and 65KO cells were rescued by expressing GRASP55-GFP or GRASP65-GFP, respectively, but not by GFP alone (lanes 4 & 7 vs. 3 & 6). (**G)** Re-expression of GRASP proteins corrects the HS and CS defects in GRASP KO cells. Confocal images of HeLa cells transfected with indicated constructs followed by HS staining. The level of HS in 55KO and 65KO cells was decreased by the expression of GRASP55-GFP or GRASP65-GFP, respectively, but not by GFP alone. Note the different HS signals in cells expressing GRASP55-or GRASP65-GFP (asterisks) vs. non-transfected cells (arrows). **(H-I)** Quantification of G. Results are presented as mean ± SEM, statistical analysis was assessed by comparing KO cells to WT cells by student’s t-test. *, p<0.05; **, p<0.01; ***, p<0.001.

To decipher the underlying mechanism of the reduced HS sulfation in GRASP KO cells, we analyzed the levels of multiple HS sulfation enzymes in the RNA-seq data and found that GRASP KO reduced the expression of the sulfur synthase PAPSS2 and the major PAPS transporter PAPST1 (SLC35B2) (Table 1). Consistently, the protein level of PAPPS2 was also significantly lower in GRASP KOs than WT cells as shown by proteomic analysis (Table 1) and Western blot (Figure 4D-E). Unlike a global reduction in HS sulfation, we only observed a significant shift from 6-sulfation (6S) to 4-sulfation (4S) in CS (Figure 3E-F). GRASP depletion largely increased the ratio of 4S/6S compared to WT. Repeating disaccharide units of CS are sulfated at C4 and C6 by C4ST-1 and C6ST-1, respectively. In our RNA-seq analysis, the mRNA levels of C6ST1 and C6ST2 were significantly decreased in GRASP KOs, especially 65KO and DKO (Table 1), which may explain the reduction of 6S in GRASP KO cells.

To confirm that the observed effects in HS and CS were caused by GRASP KO, we transfected 55KO and 65KO cells with GRASP55-GFP and GRASP65-GFP, respectively, which has previously been shown to rescue the Golgi structure and correct the defects in *N*-glycosylation and cell attachment (Xiang et al., 2013, Bekier et al., 2017, Ahat et al., 2019b). Indeed, re-expression of GRASP55 and GRASP65 in the corresponding KO cells not only restored the levels of major enzymes such as EXTL3 and GalNAcT1 (Figure 4F), but also normalized the levels of HS (Figure 4G-I) and CS (Figure S1C). Taken together, these results revealed that Golgi structure formation and defect regulate HS and CS synthesis and sulfation through modulating the expression of key enzymes.

## Discussion

In this study we found that GRASP depletion and subsequent disruption of the Golgi structure increased overall GAG synthesis but decreased their secretion. As HS and CS are the two main types of GAGs synthesized in the Golgi, we further analyzed their levels and sulfation in WT and GRASP KO cells. Our results revealed that GRASP depletion increased HS synthesis but reduced its sulfation mainly through the upregulation of EXTL3 and downregulation of PAPSS2, respectively. GRASP depletion, however, reduced CS synthesis by decreasing GalNAcT1 expression; GRASP KO also altered the balance between 4-sulfation and 6-sulfation of CS, two main forms of CS detected in LC-MS. These effects were due to GRSAP depletion as re-expression of GRASPs corrected the observed defects in HS and CS synthesis.

HS and CS levels and sulfation are tightly related to their functions in various biological processes including extracellular matrix (ECM) assembly, cell adhesion, coagulation and immune response (Aquino et al., 2010). It has been shown that abnormal sulfation of HS causes defects in FGF-2– induced proliferation and survival of multipotent progenitor cells via reducing FGF-2 and FGFR1 interaction, which contribute to Hurler syndrome (Pan et al., 2005). Similarly, in *Drosophila melanogaster* and *Caenorhabditis elegans*, reduced HS sulfation results in a delay in wound closure and defects in actin stress fiber formation (Götte et al., 2008). In another example, reduction of functional heparan sulfate proteoglycan (HSPG) has been shown to increase pericyte number while reducing its adhesion to nascent sprouts via the regulation of transforming growth factor β signal transduction (Le Jan et al., 2012). Similar to HS, CS and its synthesis enzymes are also altered in multiple disease conditions. Deficiency of an essential chondroitin synthase CHSY1 causes Temtamy preaxial brachydactyly syndrome (TPBS) (Sasarman et al., 2016). GalNAcT deficient mice showed defects in heart valve development and cardiac function via the remodulation of ECM and mitogen‑activated protein kinase (MAPK) signaling pathway (Tian et al., 2015). In developing mouse brain, the 6S level of CS is gradually decreased while 4S is gradually increased, resulting in a progressive increase of 4S/6S ratio during brain development. This change in CS sulfation was shown to reduce cortex plasticity (Miyata et al., 2012). Interestingly, in neurodegenerative diseases such as Alzheimer’s, the Golgi is fragmented possibly due to the loss of function of the GRASP proteins (Joshi et al., 2014), indicating a potential link between Golgi fragmentation, increased 4S/6S ratio, and reduced cortex plasticity in aging and neurodegenerative diseases.

How does Golgi structural defect and/or GRASP depletion affect the expression of HS and CS synthesis and sulfation enzymes is an interesting but unanswered question. This not only includes EXTL3 and GalNAcT1 that reside in the Golgi, but also PAPSS2 that is localized in the cytosol. It has been shown that many signaling molecules including mTOR, KRAS and some transcription factors such HIF1a are localized on the Golgi (Gosavi et al., 2018, Philips, 2004, Baumann et al., 2018). Golgi unstacking may affect these signaling pathways or the activity of the transcription factor, which in turn may regulate the expression of HS and CS enzymes. Similarly, it has been shown that GRASP depletion reduces the synthesis of α5β1 integrins, major cell adhesion molecules at the cell surface, which subsequently decreases cell adhesion but increases cell growth (Ahat et al., 2019b). This indicates an exciting possibility that cells may possess a sensing mechanism for Golgi structural changes, which when activated, may regulate the expression of multiple proteins to control different cellular activities.

The disruption of the Golgi stacks by GRASP-depletion used in our study is fundamentally different from the block of ER-to-Golgi trafficking by brefeldin A (BFA) treatment used in previous studies. It has been reported that disruption of the Golgi structure by BFA treatment affects HS and CS biosynthesis (Uhlin-Hansen and Yanagishita, 1993, Meneghetti et al., 2021). BFA blocks ER-to-Golgi trafficking and causes a merge of the Golgi stack (not the TGN) to the ER. Because HS and CS synthesis enzymes are localized in distinct subcompartments of the Golgi, BFA treatment affects HS and CS synthesis differently. Unlike BFA treatment, GRASP depletion disrupts the Golgi stack structure (Xiang et al, JCB 2010) but does not block membrane trafficking (Xiang et al, NatComm 2013, Ahat et al., MBoC 2019). Therefore, we believe this is the first study that evaluates the relationship of Golgi structure formation and HS/CS synthesis without blocking the flow of membrane trafficking. In addition, our systematic RNA-seq and proteomic studies identified critical genes whose alterations are responsible for the changes in HS and CS synthesis and sulfation. To our knowledge, this is the first study that links Golgi structure formation to the transcriptional regulation of O-glycosylation enzymes.

It was surprising to see that the secretion of both HS and CS was reduced in GRASP KO cells compared to WT as it has previously been shown that GRASP depletion accelerated protein trafficking through the Golgi membranes (Wang et al., 2008, Xiang et al., 2013, Lee et al., 2014). This result can be explained in several ways. First, given that all marker proteins used in the previous studies, including the vesicular stomatitis virus G (VSV-G) protein, CD8, and α5β1 integrins, are mainly modified by N-glycosylation, it is possible that GRASP depletion and/or Golgi structural disruption may affect the trafficking and secretion of different cargo molecules differently. Similarly, it has been shown that GRASP depletion alters the level of glycolipids by decreasing the level of globotriaosylceramide (Gb3) and increasing the level of monosialotetrahexosylganglioside (GM1) (Bekier et al., 2017). Second, altered sulfation of HS and CS may affect the secretion and stability of the core proteins. As sugar modifications affect protein stability and activity (Sola and Griebenow, 2009), reduced sulfation may lead to certain core proteins to be sent for degradation instead of secretion. Third, there is a possibility that more HS and CS are degraded in the conditioned media of GRASP KO cells. It has previously been shown that GRASP depletion causes missorting of lysosomal enzymes and results in their secretion (Xiang et al., 2013). It is possible that GRASP KO cells may secrete HS and CS degrading enzymes such as endoglycosidases and exohydrolases that normally reside in the lysosomes (Freeman and Hopwood, 1992). Nevertheless, the molecular mechanism that reduces HS and CS secretion in GRASP depleted cells requires further investigation.

Taken together, our results showed that disruption of the Golgi stacked structure via GRASP depletion led to the increase of total GAG synthesis, where HS level was increased due to the upregulation of EXTL3 expression and CS level was reduced because of GalNAcT1 down-regulation. In addition, Golgi defect also reduced HS sulfation via the reduction of PAPSS2. In summary, this study revealed that Golgi structural integrity and GAG synthesis are tightly linked.

## Materials and Methods

### Cell culture and transfection

Wild type, GRASP55 knockout (55KO), GRASP65 knockout (65KO), GRASP55 and GRASP65 double knockout (DKO) HeLa cells were maintained in Dulbecco’s modified Eagle’s medium (DMEM 4.5 g/l glucose) supplemented with 10% fetal bovine serum (ThermoFisher, Waltham, MA), 1% L-glutamine, 1% penicillin-streptomycin under 5% CO2 at 37°C as previously described (Bekier et al., 2017).

To express exogenous GRASP proteins, HeLa cells of ~50% confluency were transfected with indicated GRASP constructs (Xiang and Wang, 2010, Tang et al., 2010). For a 6 cm plate, 6 μg of pEGFP-N1-GRASP65 or pEGFP-N1-GRASP55 (both wild type) construct was mixed with 18 μl polyethylenimine (PEI) and 0.5 ml serum-free medium for 15 min at room temperature and then added to the cells in 4 ml DMEM containing 10% super calf serum. For control transfection, 4 μg of pEGFP-N1 construct was mixed with 18 μl PEI and 0.5 ml serum-free medium for 15 min and then added to the cells in 4 ml DMEM containing 10% super calf serum.

### Materials and sample preparation for LC-MS and PAGE analysis

Unsaturated disaccharide standards of HS (ΔUA-GlcNAc; ΔUA-GlcNS; ΔUA-GlcNAc6S; ΔUA2S-GlcNAc; ΔUA2S-GlcNS; ΔUA-GlcNS6S; ΔUA2S-GlcNAc6S; ΔUA2S-GlcNS6S), unsaturated disaccharide standards of CS (ΔUA-GalNAc; ΔUA-GalNAc4S; ΔUA-GalNAc6S; ΔUA2S-GalNAc; ΔUA2S-Gal-NAc4S; ΔUA2S-GalNAc6S; ΔUA-GalNAc4S6S; ΔUA2S-GalNAc4S6S), and unsaturated disaccharide standard of HA (ΔUA-GlcNAc), where ΔUA is 4-deoxy-α-L-threo-hex-4-enopyranosyluronic acid, were purchased from Iduron (UK). Actinase E was obtained from Kaken Biochemicals (Japan). Chondroitin lyase ABC from *Proteus vulgaris* was expressed in Linhardt’s laboratory. Recombinant Flavobacterial heparin lyases I, II, and III were expressed in Linhardt’s laboratory using *Escherichia coli* strains provided by Jian Liu (College of Pharmacy, University of North Carolina). 2-Aminoacridone (AMAC), sodium cyanoborohydride were obtained from Sigma-Aldrich (St. Louis, MO, USA). All solvents used in LC-MS were HPLC grade.

### GAG preparation for disaccharide analysis

Cells were proteolyzed at 55°C with 500 μl of 10-mg/mL actinase E for 24 h and followed by actinase E deactivation at 100°C for 30 min. The volume of the above solution containing 2 million cells was transferred to a 3-kDa molecular weight cut off (MWCO) spin tube. The filter unit was washed three times with 400 μl distilled water and then added with 300-μl digestion buffer (50 mM ammonium acetate containing 2 mM calcium chloride adjusted to pH 7.0). Recombinant heparin lyase I, II, III (pH optima 7.0−7.5) and recombinant chondroitin lyase ABC (pH optimum 7.4, 10 mU each) were added to each filter unit containing sample and mixed well. The samples were all incubated at 37 °C for 24 h. The enzymatic digestion was terminated by ultrafiltration through the 3-kDa spin tube. The filtrate was collected, and the filter unit was washed twice with 200 μl distilled water. All the filtrates containing the disaccharide products were combined and dried via freeze dry. For the medium samples, 400-μl medium from each specimen was ultrafiltrated through a 3-kDa molecular weight cut off (MWCO) spin tube to remove small molecular compounds, and then went through the same digestion procedure as mentioned above.

### AMAC labeling of disaccharides

The dried samples were AMAC-labeled by adding 10 μl of 0.1 M AMAC in DMSO/acetic acid (17/3, V/V) incubating at room temperature for 10 min, followed by adding 10 μl of 1 M aqueous sodium cyanoborohydride and incubating for 1 h at 45°C.A mixture containing all 17-disaccharide standards prepared at 0.5ng/μl was similarly AMAC-labeled and used for each run as an external standard. After the AMAC-labeling reaction, the samples were centrifuged, and each supernatant was recovered.

### LC-MS

LC was performed on an Agilent 1200 LC system at 45 °C using an Agilent Poroshell 120 ECC18 (2.7 μm, 3.0 × 50 mm) column. Mobile phase A (MPA) was 50 mM ammonium acetate aqueous solution, and the mobile phase B (MPB) was methanol. The mobile phase passed through the column at a flow rate of 300 μl/min. The gradient was 0-10 min, 5-45% B; 10-10.2 min, 45-100%B; 10.2-14 min, 100%B; 14-22 min, 100-5%B. Injection volume is 5 μl.

A triple quadrupole mass spectrometry system equipped with an ESI source (Thermo Fisher Scientific, San Jose, CA) was used a detector. The online MS analysis was at the Multiple Reaction Monitoring (MRM) mode. MS parameters: negative ionization mode with a spray voltage of 3000 V, a vaporizer temperature of 300°C, and a capillary temperature of 270°C.

### Western blot

Wild type and GRASP KO cells are lysed in 20 mM Tris-HCl, pH 8.0, 150 mM NaCl, 1% Triton X-100 and protease inhibitors for 30 min on ice. Lysates were cleared by centrifugation (20,000 *g* for 20 min at 4°C). After electrophoresis and transfer, nitrocellular membranes were incubated with antibodies to actin (Sigma, A2066), EXT1 (Santa Cruz, sc-515144), EXT2 (Santa Cruz, sc-514092), EXTL3 (Santa Cruz, sc-271986), GalNAcT1 (Novus, NBP1-81852), GFP (Proteintech, 66002-1-Ig), GRASP55 (Proteintech, 10598-1-AP), GRASP65 (Santa Cruz, sc-374423), or PAPSS2 (Santa Cruz, sc-271429) overnight at 4°C. The membranes were extensively washed and further incubated with HRP conjugated goat anti-Rabbit or goat anti-mouse secondary antibodies for 1 h at room temperature and exposed to a FluorChem M machine (Proteinsimple).

### Immunofluorescence microscopy

Cells were grown on sterile glass coverslips and rinsed with phosphate buffered saline (PBS) before fixation. For total protein staining, cells were fixed in 4% paraformaldehyde for 10 min and permeabilized with 0.2% Triton X-100 in PBS for 10 min. For cell surface staining, cells were fixed in 1% paraformaldehyde for 10 min and not permeabilized. Cells were incubated with primary antibodies for HS (10E4, Amsbio 370255, 1:100) and CS (CS-56, Abcam ab11570, 1:50) overnight at 4°C, washed and probed with the appropriate secondary antibodies conjugated to TRITC for 45 min. To confirm that the detected 10E4 signal was specific to HS, the HS antibody (10E4 1:100) was incubated with or without 40 μg/ml heparin overnight at 4°C and then used for primary antibody staining of fixed cells. DNA was stained with Hoechst for 5 min. Coverslips were rinsed with PBS and mounted with Mowiol onto slides. Images were taken with a 20x air objective or a 63x oil objective on a Nikon ECLIPSE Ti2 Confocal microscope and shown as max projections.

### Flow cytometry

HeLa cells (WT and KO) were detached using 20 mM EDTA and resuspended in PBS with 0.5% BSA. The cells were fixed and permeabilized with 4% PFA for 10 min and 0.2% Triton X-100 for 10 min, respectively. After washing with PBS twice, the cells were incubated with the primary antibody for HS (10E4, 1:100) or CS (CS-56, 1:50) (or control without a primary antibody) with rotation for 1.5 h at room temperature. Both primary antibodies and the control group were incubated with goat anti mouse secondary antibodies (TRITC) for 1 h with rotation at room temperature. The cells were sorted with a Sony MA900 Multi-Application Cell Sorter and the data was analyzed with FlowJo software.

### Proteomics analysis

#### Sample Preparation

Three replicates of each WT, 55KO and 65KO cells were propagated as described above in 15 cm dishes. Upon achieving 80% confluency, the growth media were aspirated, and the cells were washed with PBS five times, then changed to 20 ml serum free medium and further incubated for 12 h. The media were first cleared by centrifugation at 500 *g* for 10 min at 4°C and then at 4000 *g* for 15 min at 4°C, and then filtered with a 0.45 μm filter. The cleaned media were concentrated with a 3 kDa cutoff ultrafilter (Millipore, UFC900324) to 200-300 μl, and the protein concentration was determined with Bradford assay (Bio-Rad, Cat # 5000006). For cell lysates collection, after removing the media, cells were washed with PBS twice, and collected in 10 ml PBS by scraping, lysed in Pierce™ RIPA buffer (Thermo, 89900) with a protein inhibitor cocktail (Thermo). The protein concentration was tested with Bradford assay and normalized, 75 μg of each sample was provided to the Mass Spectrometry-Based Proteomics Resource Facility at Department of Pathology, University of Michigan for TMT labeling, LC-MS/MS and bioinformatics analysis.

#### Protein Digestion and TMT labeling

Samples were proteolysed and labeled with TMT 10-plex essentially by following manufacturer’s protocol (ThermoFisher, Cat # 90110, Lot # VJ306782). Briefly, upon reduction and alkylation of cysteines, the proteins were precipitated by adding 6 volumes of ice-cold acetone followed by overnight incubation at −20°C. The precipitate was spun down, and the pellet was allowed to air dry. The pellet was resuspended in 0.1 M TEAB and overnight digestion with trypsin (1:50; enzyme:protein) at 37°C was performed with constant mixing using a thermomixer. The TMT 10-plex reagents were dissolved in 41 μl of anhydrous acetonitrile and labeling was performed by transferring the entire digest to TMT reagent vial and incubating at room temperature for 1 h. Reaction was quenched by adding 8 μl of 5% hydroxyl amine and further 15 min incubation. Labeled samples were mixed, and dried using a vacufuge. An offline fractionation of the combined sample (~200 μg) into 8 fractions was performed using high pH reversed-phase peptide fractionation kit according to the manufacturer’s protocol (Pierce; Cat # 84868). Fractions were dried and reconstituted in 9 μl of 0.1% formic acid/2% acetonitrile in preparation for LC-MS/MS analysis.

#### Liquid chromatography-mass spectrometry analysis (LC-multinotch MS3)

In order to obtain superior quantitation accuracy, we employed multinotch-MS3 (McAlister GC) which minimizes the reporter ion ratio distortion resulting from fragmentation of co-isolated peptides during MS analysis. Orbitrap Fusion (Thermo Fisher Scientific) and RSLC Ultimate 3000 nano-UPLC (Dionex) was used to acquire the data. 2 μl of the sample was resolved on a PepMap RSLC C18 column (75 μm i.d. × 50 cm; Thermo Scientific) at the flowrate of 300 nl/min using 0.1% formic acid/acetonitrile gradient system (2-22% acetonitrile in 150 min;22-32% acetonitrile in 40 min; 20 min wash at 90% followed by 50 min re-equilibration) and directly spray onto the mass spectrometer using EasySpray source (Thermo Fisher Scientific). Mass spectrometer was set to collect one MS1 scan (Orbitrap; 120K resolution; AGC target 2×105; max IT 100 ms) followed by data-dependent, “Top Speed” (3 seconds) MS2 scans (collision induced dissociation; ion trap; NCE 35; AGC 5×103; max IT 100 ms). For multinotch-MS3, top 10 precursors from each MS2 were fragmented by HCD followed by Orbitrap analysis (NCE 55; 60K resolution; AGC 5×104; max IT 120 ms, 100-500 m/z scan range).

#### Data analysis

Proteome Discoverer (v2.4; Thermo Fisher) was used for data analysis. MS2 spectra were searched against SwissProt human protein database (20353 entries; downloaded on 06/20/2019) using the following search parameters: MS1 and MS2 tolerance were set to 10 ppm and 0.6 Da, respectively; carbamidomethylation of cysteines (57.02146 Da) and TMT labeling of lysine and N-termini of peptides (229.16293 Da) were considered static modifications; oxidation of methionine (15.9949 Da) and deamidation of asparagine and glutamine (0.98401 Da) were considered variable. Identified proteins and peptides were filtered to retain only those that passed ≤1% FDR threshold. Quantitation was performed using high-quality MS3 spectra (Average signal-to-noise ratio of 10 and <50% isolation interference).

### RNA-Seq analysis

RNA samples were collected from each of the four HeLa cell lines: WT, 55KO, 65KO, and DKO. Cells at an exponential growth phase (~ 80% confluency) in 6 well dishes were collected. Five replicates of each cell line were lysed using Trizol. RNA samples were prepared using the Direct-zol™ RNA Miniprep Plus kit and treated with DNase I provided in the same kit. The samples were sent to UMich Advanced Genomic Core for library creation and processing. At the core, after passing quality control for quantity and purity, RNA samples were used to create 3’ mRNA libraries using the QuantSeq 3’ mRNA-Seq Library Prep Kit FWD for Illumina kit with the UMI add-on kit (Lexogen, Cat # 081.96), which used oligoT priming to generate the first cDNA strand from RNAs with a poly-A tail. Single-read sequencing for the cDNA library was performed on Illumina NextSeq sequencer for 100 cycles. Sequencing results were trimmed using Trim Galore (v 0.5.0). Alignment of reads to human genome GRCh38 from ENSEMBL (https://useast.ensembl.org/index.html) was performed in house using STAR (v 2.6.0). Trim_Galore and STAR can be found on https://www.bioinformatics.babraham.ac.uk/projects/trim_galore/ and https://www.ncbi.nlm.nih.gov/pmc/articles/PMC3530905/, respectively.

### Quantification and statistics

In all figures, the quantification results are expressed as the mean ± SEM (standard error of the mean) from 3-5 independent experiments, unless otherwise stated. The statistical significance of the results was assessed using Student’s *t*-test. *, *p<*0.05, **, *p*<0.01, ***, *p*<0.001.

## Supplemental Information

Supplemental Information includes one supplemental figure and six supplemental tables.

## Acknowledgments

We thank Ge Yu, Judith Meyers Opp and the University of Michigan Advanced Genomics Core for their contribution in the RNA-Seq experiment. We thank Drs. Venkatesha Basrur, Felipe da Veiga Leprevost, Alexey Nesvizhskii and the Mass Spectrometry-Based Proteomics Resource Facility in the Department of Pathology at the University of Michigan for their contribution in Proteomics analysis. We thank the members of the Wang lab and Linhardt lab for stimulating discussions and technical support. This work was supported by the National Institutes of Health (Grant R35GM130331), Mizutani Foundation for Glycoscience, MCubed, and the Fast Forward Protein Folding Disease Initiative of the University of Michigan to Y. Wang, and a University of Michigan Rackham Predoctoral fellowship to E. Ahat.

## Author Contributions

E.A., Y.S., F.Z., R.L. and Y.W. designed the experiments; E.A. prepared the samples and Y.S., K.X. performed labeling and LC-MS experiments. E.A. performed most of the other experiments. W.R. and S.B. performed data search and analysis from the RNA-seq and Proteomics experiments. J.L. and E.A. performed the flow cytometry experiment. E.A., Y.S., R.L. and Y.W. analyzed the data. E.A. and Y.W. made the figures and wrote the first draft; Y.S. K.X., F.Z. and R.L. edited the manuscript. All authors discussed the results and contributed to the final manuscript.

## Supplemental information

**Supplemental Figure 1.**
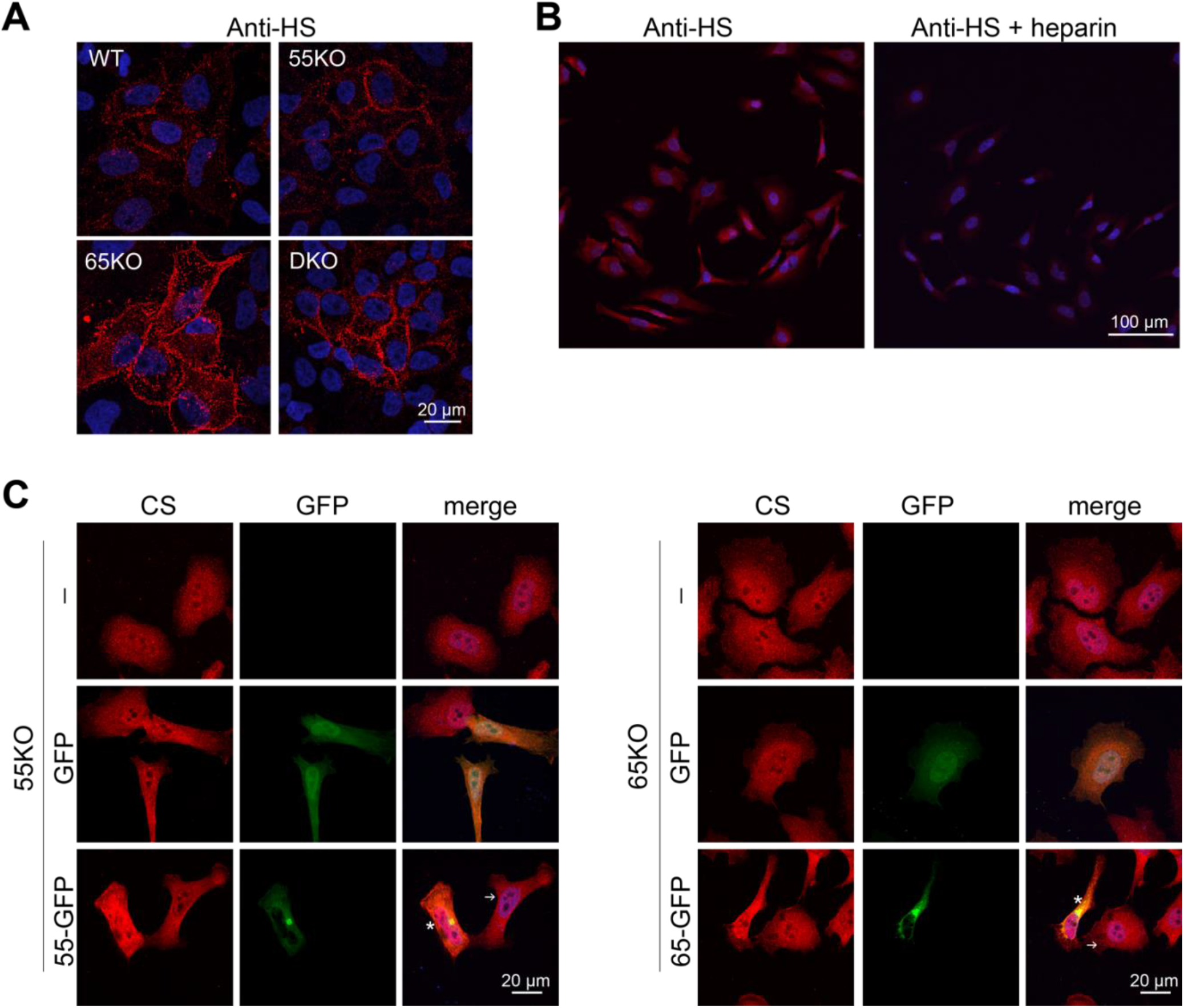
GRASP KO increases HS level at the cell surface and GRASP re-expression rescues CS defects in GRASP KO cells. **(A)** GRASP KO increases HS level at the cell surface. Confocal images of indicated cell lines stained with an anti-HS antibody (10E4) without permeabilization. **(B)** The HS signal of the anti-HS antibody is specific. Permeabilized WT cells were stained with the anti-HS antibody (left panel) or with the anti-HS antibody preincubated with excess amount of heparin (right) before imaging. **(C)** Rescue of the Golgi structure by GRASP re-expression corrects the CS defects in GRASP KO cells. Confocal images of 55KO or 65KO cells transfected with indicated constructs and stained for CS with the CS-56 antibody. Scale bars in all panels, 20 μm.

**Supplemental Table 1.** The amount of three kinds of GAG from cell samples. Cell lysate from indicated cells were analyzed for total GAGs by LC-MS, results are normalized to one million cells. Five technical replicates are shown for each cell line.

| Sample |  | Total cell<br>number<br>(million) | Amount of GAG<br>(ng) per million<br>cells | Amount of each kind of GAG (ng) per<br>million cells |  |  |
| --- | --- | --- | --- | --- | --- | --- |
|  |  |  |  | HS | CS | HA |
| WT | 1 | 7.8 | 319.21 | 194.28 | 118.17 | 6.76 |
|  | 2 | 7.5 | 330.74 | 202.97 | 121.16 | 6.61 |
|  | 3 | 8.0 | 350.40 | 208.69 | 134.50 | 7.21 |
|  | 4 | 8.5 | 348.29 | 209.07 | 131.77 | 7.45 |
|  | 5 | 7.8 | 318.55 | 192.44 | 119.37 | 6.74 |
| G55KO | 1 | 13.0 | 383.04 | 251.87 | 117.20 | 13.97 |
|  | 2 | 12.5 | 400.88 | 260.51 | 126.35 | 14.02 |
|  | 3 | 12.1 | 394.46 | 256.32 | 123.61 | 14.53 |
|  | 4 | 11.9 | 443.26 | 292.82 | 135.39 | 15.06 |
|  | 5 | 12.0 | 399.19 | 263.36 | 122.38 | 13.45 |
| G65KO | 1 | 12.2 | 524.01 | 388.18 | 122.26 | 13.57 |
|  | 2 | 10.5 | 585.35 | 425.57 | 144.65 | 15.12 |
|  | 3 | 11.7 | 522.45 | 381.23 | 127.40 | 13.81 |
|  | 4 | 10.9 | 569.16 | 426.28 | 128.02 | 14.85 |
|  | 5 | 11.7 | 517.92 | 379.43 | 124.36 | 14.13 |
| DKO | 1 | 12.5 | 464.67 | 295.77 | 149.16 | 19.74 |
|  | 2 | 12.9 | 442.95 | 286.77 | 138.89 | 17.29 |
|  | 3 | 12.6 | 458.21 | 293.83 | 146.51 | 17.87 |
|  | 4 | 13.4 | 463.56 | 290.90 | 152.24 | 20.41 |
|  | 5 | 14.0 | 414.50 | 267.46 | 128.44 | 18.59 |

**Supplemental Table 2.** The amount of three kinds of GAG from medium samples. Conditioned media from indicated cells were analyzed for total GAGs by LC-MS. Five technical replicates are shown for each cell line. GAG amounts per million cells were calculated and shown in Figure 1D.

| Sample |  | Total cell number (million) | Total amount of GAG in the medium (ng) | Amount of each kind of GAG (ng) |  |  |
| --- | --- | --- | --- | --- | --- | --- |
|  |  |  |  | HS | CS | HA |
| WT-M | 1 | 7.8 | 1182.12 | 267.40 | 436.71 | 478.01 |
|  | 2 | 7.5 | 1246.76 | 362.17 | 448.02 | 436.57 |
|  | 3 | 8.0 | 1356.15 | 368.26 | 502.05 | 485.84 |
|  | 4 | 8.5 | 1011.04 | 370.29 | 369.62 | 271.13 |
|  | 5 | 7.8 | 1179.66 | 345.07 | 390.76 | 443.83 |
| G55KO-M | 1 | 13.0 | 1381.84 | 540.55 | 256.11 | 585.18 |
|  | 2 | 12.5 | 1098.50 | 493.11 | 157.56 | 447.83 |
|  | 3 | 12.1 | 1358.76 | 545.61 | 262.69 | 550.46 |
|  | 4 | 11.9 | 1366.89 | 684.66 | 203.67 | 478.55 |
|  | 5 | 12.0 | 1405.03 | 561.53 | 288.87 | 554.62 |
| G65KO-M | 1 | 12.2 | 1251.66 | 537.08 | 235.84 | 478.73 |
|  | 2 | 10.5 | 1288.60 | 584.23 | 253.50 | 450.86 |
|  | 3 | 11.7 | 1345.90 | 597.45 | 276.86 | 471.59 |
|  | 4 | 10.9 | 1337.10 | 532.68 | 250.96 | 553.46 |
|  | 5 | 11.7 | 1104.74 | 491.10 | 190.63 | 423.01 |
| DKO-M | 1 | 12.5 | 1332.16 | 492.36 | 320.53 | 519.27 |
|  | 2 | 12.9 | 1308.39 | 459.76 | 337.71 | 510.92 |
|  | 3 | 12.6 | 1441.38 | 489.11 | 374.16 | 578.11 |
|  | 4 | 13.4 | 1323.26 | 431.75 | 366.56 | 524.95 |
|  | 5 | 14.0 | 1320.19 | 439.70 | 232.61 | 647.88 |

**Supplemental Table 3.** The percentage of each detected HS disaccharide from cell samples. Cell lysate from indicated cells were analyzed for HS by LC-MS, results are normalized to one million cells. Five technical replicates are shown for each cell line.

| Sample |  | HS disaccharides (%) |  |  |  |  |  |  |  |
| --- | --- | --- | --- | --- | --- | --- | --- | --- | --- |
|  |  | TriS | NS6S | NS2S | NS | 2S6S | 6S | 2S | 0S |
| WT | 1 | 3.06 | 6.37 | 3.02 | 14.31 | 0.28 | 12.72 | 0.66 | 59.57 |
|  | 2 | 3.11 | 6.36 | 3.09 | 13.55 | 0.28 | 12.63 | 0.67 | 60.30 |
|  | 3 | 3.13 | 6.30 | 2.89 | 14.33 | 0.29 | 12.39 | 0.39 | 60.28 |
|  | 4 | 2.84 | 6.21 | 2.91 | 14.33 | 0.29 | 12.52 | 0.64 | 60.28 |
|  | 5 | 3.04 | 6.36 | 3.03 | 15.03 | 0.30 | 12.61 | 0.43 | 59.20 |
| G55KO | 1 | 2.11 | 5.50 | 2.46 | 11.68 | 0.23 | 11.22 | 0.35 | 66.45 |
|  | 2 | 2.39 | 5.82 | 2.70 | 12.09 | 0.26 | 11.67 | 0.46 | 64.62 |
|  | 3 | 2.02 | 5.52 | 2.64 | 12.21 | 0.19 | 11.76 | 0.42 | 65.24 |
|  | 4 | 2.03 | 5.51 | 2.57 | 12.16 | 0.25 | 11.26 | 0.53 | 65.69 |
|  | 5 | 2.05 | 5.46 | 2.68 | 12.21 | 0.22 | 11.17 | 0.43 | 65.79 |
| G65KO | 1 | 2.14 | 4.64 | 2.61 | 12.77 | 0.23 | 10.83 | 0.83 | 65.97 |
|  | 2 | 2.03 | 4.46 | 2.48 | 12.62 | 0.23 | 11.05 | 0.95 | 66.19 |
|  | 3 | 2.18 | 4.50 | 2.44 | 12.61 | 0.23 | 10.86 | 0.76 | 66.41 |
|  | 4 | 2.05 | 4.40 | 2.30 | 12.56 | 0.23 | 10.08 | 1.01 | 67.38 |
|  | 5 | 2.16 | 4.55 | 2.41 | 12.94 | 0.25 | 10.65 | 0.80 | 66.24 |
| DKO | 1 | 1.72 | 4.50 | 1.51 | 11.15 | 0.24 | 10.71 | 0.51 | 69.65 |
|  | 2 | 1.81 | 4.63 | 1.46 | 10.94 | 0.26 | 10.66 | 0.52 | 69.73 |
|  | 3 | 1.72 | 4.68 | 1.42 | 11.31 | 0.24 | 10.84 | 0.63 | 69.16 |
|  | 4 | 1.75 | 4.60 | 1.46 | 10.89 | 0.24 | 10.89 | 0.43 | 69.74 |
|  | 5 | 1.50 | 4.51 | 1.41 | 10.65 | 0.24 | 10.49 | 0.51 | 70.69 |

**Supplemental Table 4.** The percentage of each detected HS disaccharide from medium samples. Conditioned media from indicated cells were analyzed for HS by LC-MS. Five technical replicates are shown for each cell line.

| Sample |  | HS disaccharides (%) |  |  |  |  |  |  |  |
| --- | --- | --- | --- | --- | --- | --- | --- | --- | --- |
|  |  | TriS | NS6S | NS2S | NS | 2S6S | 6S | 2S | 0S |
| WT-M | 1 | 0.17 | 5.91 | 2.62 | 14.48 | 0.66 | 7.68 | 0.14 | 68.35 |
|  | 2 | 0.19 | 6.55 | 2.20 | 15.61 | 0.10 | 7.13 | 0.01 | 68.21 |
|  | 3 | 0.19 | 6.22 | 1.52 | 13.04 | 0.20 | 11.53 | 0.01 | 67.30 |
|  | 4 | 0.11 | 5.38 | 1.54 | 13.96 | 0.00 | 8.87 | 0.15 | 69.98 |
|  | 5 | 0.25 | 7.51 | 1.97 | 14.62 | 0.12 | 12.16 | 0.01 | 63.36 |
| G55KO-M | 1 | 0.33 | 7.29 | 1.76 | 15.27 | 0.08 | 4.68 | 0.00 | 70.58 |
|  | 2 | 0.17 | 6.97 | 1.17 | 17.13 | 0.17 | 6.02 | 0.00 | 68.37 |
|  | 3 | 0.32 | 6.46 | 2.14 | 14.95 | 0.27 | 5.77 | 0.07 | 70.02 |
|  | 4 | 0.51 | 6.80 | 1.93 | 16.23 | 0.17 | 8.39 | 0.00 | 65.98 |
|  | 5 | 0.18 | 7.61 | 1.78 | 13.62 | 0.09 | 6.84 | 0.00 | 69.89 |
| G65KO-M | 1 | 0.37 | 7.50 | 2.50 | 15.82 | 0.19 | 6.45 | 0.00 | 67.15 |
|  | 2 | 0.36 | 7.60 | 1.46 | 15.20 | 0.28 | 9.28 | 0.18 | 65.62 |
|  | 3 | 0.30 | 7.47 | 2.84 | 16.26 | 0.33 | 7.28 | 0.11 | 65.40 |
|  | 4 | 0.00 | 5.94 | 1.71 | 11.87 | 0.14 | 6.57 | 0.00 | 73.76 |
|  | 5 | 0.17 | 6.33 | 2.15 | 16.54 | 0.19 | 7.23 | 0.00 | 67.39 |
| DKO-M | 1 | 0.19 | 6.42 | 3.39 | 10.37 | 0.60 | 3.21 | 0.01 | 75.82 |
|  | 2 | 0.00 | 5.37 | 1.84 | 8.90 | 0.29 | 6.48 | 0.00 | 77.12 |
|  | 3 | 0.17 | 9.08 | 2.17 | 8.40 | 2.42 | 3.77 | 0.01 | 73.98 |
|  | 4 | 0.23 | 7.99 | 1.07 | 9.64 | 0.21 | 7.99 | 0.00 | 72.86 |
|  | 5 | 0.27 | 7.70 | 2.00 | 12.44 | 0.30 | 6.26 | 0.00 | 71.02 |

**Supplemental Table 5.** The percentage of each detected CS disaccharide from cell samples. Cell lysate from indicated cells were analyzed for CS by LC-MS, results are normalized to one million cells. Five technical replicates are shown for each cell line.

| Sample |  | CS disaccharides (%) |  |  |  |  |  |  |  |
| --- | --- | --- | --- | --- | --- | --- | --- | --- | --- |
|  |  | TriS | 2S4S | 2S6S | 4S6S | 4S | 6S | 2S | 0S |
| WT | 1 | 0.03 | 0.30 | 2.31 | 2.05 | 75.28 | 18.87 | 0.00 | 1.14 |
|  | 2 | 0.01 | 0.38 | 2.09 | 1.97 | 78.45 | 16.10 | 0.00 | 1.00 |
|  | 3 | 0.02 | 0.37 | 2.05 | 1.90 | 78.87 | 15.76 | 0.00 | 1.03 |
|  | 4 | 0.02 | 0.38 | 1.73 | 2.12 | 78.49 | 16.13 | 0.00 | 1.13 |
|  | 5 | 0.02 | 0.24 | 1.92 | 2.06 | 73.82 | 21.10 | 0.00 | 0.84 |
| G55KO | 1 | 0.13 | 0.76 | 2.38 | 1.65 | 81.11 | 13.32 | 0.00 | 0.66 |
|  | 2 | 0.04 | 0.80 | 2.80 | 1.87 | 78.89 | 14.84 | 0.00 | 0.75 |
|  | 3 | 0.05 | 0.77 | 2.75 | 1.78 | 78.35 | 15.44 | 0.00 | 0.86 |
|  | 4 | 0.04 | 0.83 | 2.49 | 1.54 | 79.13 | 14.87 | 0.00 | 1.11 |
|  | 5 | 0.07 | 0.81 | 2.68 | 1.87 | 79.58 | 14.22 | 0.00 | 0.77 |
| G65KO | 1 | 0.01 | 0.38 | 1.13 | 1.72 | 87.58 | 8.63 | 0.00 | 0.55 |
|  | 2 | 0.01 | 0.23 | 1.25 | 1.70 | 85.73 | 10.32 | 0.00 | 0.76 |
|  | 3 | 0.02 | 0.31 | 1.36 | 1.59 | 84.01 | 11.98 | 0.00 | 0.72 |
|  | 4 | 0.01 | 0.38 | 1.29 | 1.63 | 87.37 | 8.71 | 0.00 | 0.60 |
|  | 5 | 0.04 | 0.36 | 1.12 | 1.64 | 86.05 | 10.16 | 0.00 | 0.64 |
| DKO | 1 | 0.08 | 0.97 | 1.74 | 1.87 | 84.94 | 9.65 | 0.00 | 0.75 |
|  | 2 | 0.05 | 0.73 | 1.73 | 1.76 | 86.61 | 8.71 | 0.00 | 0.40 |
|  | 3 | 0.04 | 0.85 | 1.76 | 1.88 | 85.79 | 9.06 | 0.00 | 0.63 |
|  | 4 | 0.07 | 0.79 | 1.70 | 1.87 | 85.39 | 9.53 | 0.00 | 0.66 |
|  | 5 | 0.08 | 0.90 | 1.77 | 1.90 | 85.56 | 8.76 | 0.00 | 1.03 |

**Supplemental Table 6.** The percentage of each detected CS disaccharide from medium samples. Conditioned media from indicated cells were analyzed for CS by LC-MS. Five technical replicates are shown for each cell line.

| Sample |  | CS disaccharides (%) |  |  |  |  |  |  |  |
| --- | --- | --- | --- | --- | --- | --- | --- | --- | --- |
|  |  | TriS | 2S4S | 2S6S | 4S6S | 4S | 6S | 2S | 0S |
| WT-M | 1 | 0.12 | 0.00 | 2.22 | 0.78 | 75.42 | 20.04 | 0.01 | 1.40 |
|  | 2 | 0.00 | 0.78 | 0.94 | 1.28 | 83.32 | 13.68 | 0.01 | 0.00 |
|  | 3 | 0.00 | 0.35 | 1.00 | 0.96 | 82.93 | 14.40 | 0.00 | 0.36 |
|  | 4 | 0.00 | 0.71 | 2.56 | 1.29 | 82.27 | 13.16 | 0.01 | 0.00 |
|  | 5 | 0.00 | 0.22 | 2.19 | 1.63 | 79.71 | 16.25 | 0.01 | 0.00 |
| G55KO-M | 1 | 0.00 | 1.09 | 2.64 | 1.90 | 77.56 | 16.53 | 0.01 | 0.27 |
|  | 2 | 0.00 | 1.31 | 2.15 | 1.09 | 73.76 | 21.66 | 0.01 | 0.00 |
|  | 3 | 0.00 | 1.27 | 1.85 | 0.90 | 69.86 | 17.63 | 0.01 | 8.49 |
|  | 4 | 0.00 | 0.60 | 3.36 | 0.87 | 74.88 | 18.56 | 0.01 | 1.72 |
|  | 5 | 0.00 | 0.00 | 4.22 | 1.62 | 72.77 | 18.60 | 0.01 | 2.79 |
| G65KO-M | 1 | 0.00 | 0.35 | 2.40 | 0.71 | 78.54 | 8.66 | 0.01 | 9.33 |
|  | 2 | 0.00 | 0.41 | 1.01 | 0.67 | 86.18 | 9.38 | 0.01 | 2.34 |
|  | 3 | 0.00 | 0.72 | 0.97 | 0.76 | 88.41 | 8.36 | 0.01 | 0.77 |
|  | 4 | 0.00 | 0.25 | 0.86 | 1.03 | 83.47 | 8.69 | 0.01 | 5.69 |
|  | 5 | 0.00 | 0.73 | 0.35 | 0.90 | 82.55 | 10.16 | 0.01 | 5.30 |
| DKO-M | 1 | 0.00 | 1.36 | 1.81 | 0.67 | 87.24 | 8.91 | 0.01 | 0.00 |
|  | 2 | 0.17 | 0.10 | 0.94 | 1.44 | 90.08 | 6.72 | 0.01 | 0.54 |
|  | 3 | 0.00 | 0.14 | 1.51 | 0.70 | 86.15 | 8.96 | 0.01 | 2.53 |
|  | 4 | 0.00 | 0.77 | 1.10 | 1.31 | 88.57 | 6.86 | 0.01 | 1.38 |
|  | 5 | 0.00 | 1.94 | 2.29 | 1.06 | 85.54 | 9.02 | 0.01 | 0.14 |

